# IRIS-FGM: an integrative single-cell RNA-Seq interpretation system for functional gene module analysis

**DOI:** 10.1101/2020.11.04.369108

**Authors:** Yuzhou Chang, Carter Allen, Changlin Wan, Dongjun Chung, Chi Zhang, Zihai Li, Qin Ma

## Abstract

**Summary:** Single-cell RNA-Seq (scRNA-Seq) data is useful in discovering cell heterogeneity and signature genes in specific cell populations in cancer and other complex diseases. Specifically, the investigation of functional gene modules (FGM) can help to understand gene interactive networks and complex biological processes. QUBIC2 is recognized as one of the most efficient and effective tools for FGM identification from scRNA-Seq data. However, its limited availability to a C implementation restricted its application to only a few downstream analyses functionalities. We developed an R package named IRIS-FGM (Integrative scRNA-Seq Interpretation System for Functional Gene Module analysis) to support the investigation of FGMs and cell clustering using scRNA-Seq data. Empowered by QUBIC2, IRIS-FGM can effectively identify co-expressed and co-regulated FGMs, predict cell types/clusters, uncover differentially expressed genes, and perform functional enrichment analysis. It is noteworthy that IRIS-FGM can also takes Seurat objects as input, which facilitate easy integration with existing analysis pipeline.

**Availability and Implementation:** IRIS-FGM is implemented in R environment (as of version 3.6) with the source code freely available at https://github.com/OSU-BMBL/IRIS-FGM

**Supplementary information:** Supplementary data are available at *Bioinformatics* online.

## 1 Introduction

Single-cell RNA-Seq (scRNA-Seq) data characterizes the cell heterogeneity in complex tissues and diseases that can reveal cell subpopulations and their unique gene expression patterns. Biclustering is a widely accepted approach for identifying co-expressed genes under subsets of cells in a gene expression dataset. Our previously developed tool, QUBIC2 (Xie, et al., 2019), outperformed existing methods, such as FABIA (Hochreiter, et al., 2010), ISA (Bergmann, et al., 2003), Plaid (Lazzeroni and Owen, 2002), and Bimax (Prelic, et al., 2006) in identifying biologically meaningful biclusters on 10X scRNA-Seq data and was successfully used for cell-type-specific regulon prediction, which revealed regulatory signals and their targeted gene in a specific cell type (Ma, et al., 2020). Using a left-truncated mixture Gaussian (LTMG) model (Wan, et al., 2019), it identifies biclusters, genes within which are simultaneously co-expressed and co-regulated, i.e., a functional gene module (FGM). The investigation of FGM can help to understand gene-gene interaction networks and complex biological processes from scRNA-Seq data. However, previously QUBIC2 was only available as a C implementation, and its applicative power was also restricted to only a few downstream analyses functionalities. Furthermore, because of success in our comprehensive web server-based RNA-Seq interpretation system (Monier, et al., 2019), a powerful and multiple functional interpretation system will improve usability and interpretability of analysis. To the end, we developed an R package named IRIS-FGM (<u>I</u>ntegrative sc<u>R</u>NA-Seq <u>I</u>nterpretation <u>S</u>ystem for <u>F</u>unctional <u>G</u>ene <u>M</u>odule analysis) to support the investigation of FGMs and cell clustering using scRNA-Seq data. Empowered by QUBIC2, IRIS-FGM can effectively identify co-expressed and co-regulated FGMs, predict cell types/clusters, uncover differentially expressed gene patterns, and perform functional enrichment analysis.

The IRIS-FGM framework consists of three key steps (Figure 1A). In the first step, the raw scRNA-Seq data is read as a Seurat object which is preprocessed by normalizing expression values, as well as removing low-quality genes and cells. The LTMG is then applied to deconvolve the normalized read counts into multiple signal components and generate the discretized gene-cell matrix. In the second step, the QUBIC2 algorithm is applied to identify biclusters (i.e., co-expressed and co-regulated FGMs) from the LTMG discretized matrix and generate a cell-cell distance matrix/cell-cell graph by combining all biclusters. In such a graph, each node represents a cell and each edge represents the occurrence of the connected two cells within the same bicluster (Xie, et al., 2019). We further identify cell clusters based on the cell-cell graph by using the Markov clustering algorithm (MCL) and implement the Seurat *FindMarkers* function for differential gene analysis (Butler, et al., 2018). In a comparison study of cell clustering approaches on the test dataset, IRIS-FGM outperforms other five popular biclustering tools (QUBIC (Li, et al., 2009), FABIA, ISA, Plaid, Bimax) and four clustering tools (i.e., SC3 (Kiselev, et al., 2017), SINCERA (Guo, et al., 2015), SNN-Cliq (Shi and Huang, 2017), and Seurat (Butler, et al., 2018)) (Figure 1B, **Supplementary Method.S1**). IRIS-FGM contains multiple visualization functions that allows users to visualize the analytical results of FGM and cell clustering results, such as UMAP plot, FGM heatmap (Figure 1C), and FGM network (Figure 1D). Detailed codes can be found in the next section. As IRIS-FGM uses Seurat object, Seurat clustering results from raw expression matrix or LTMG discretized matrix can also be directly fed into IRIS-FGM.

**Figure 1.**
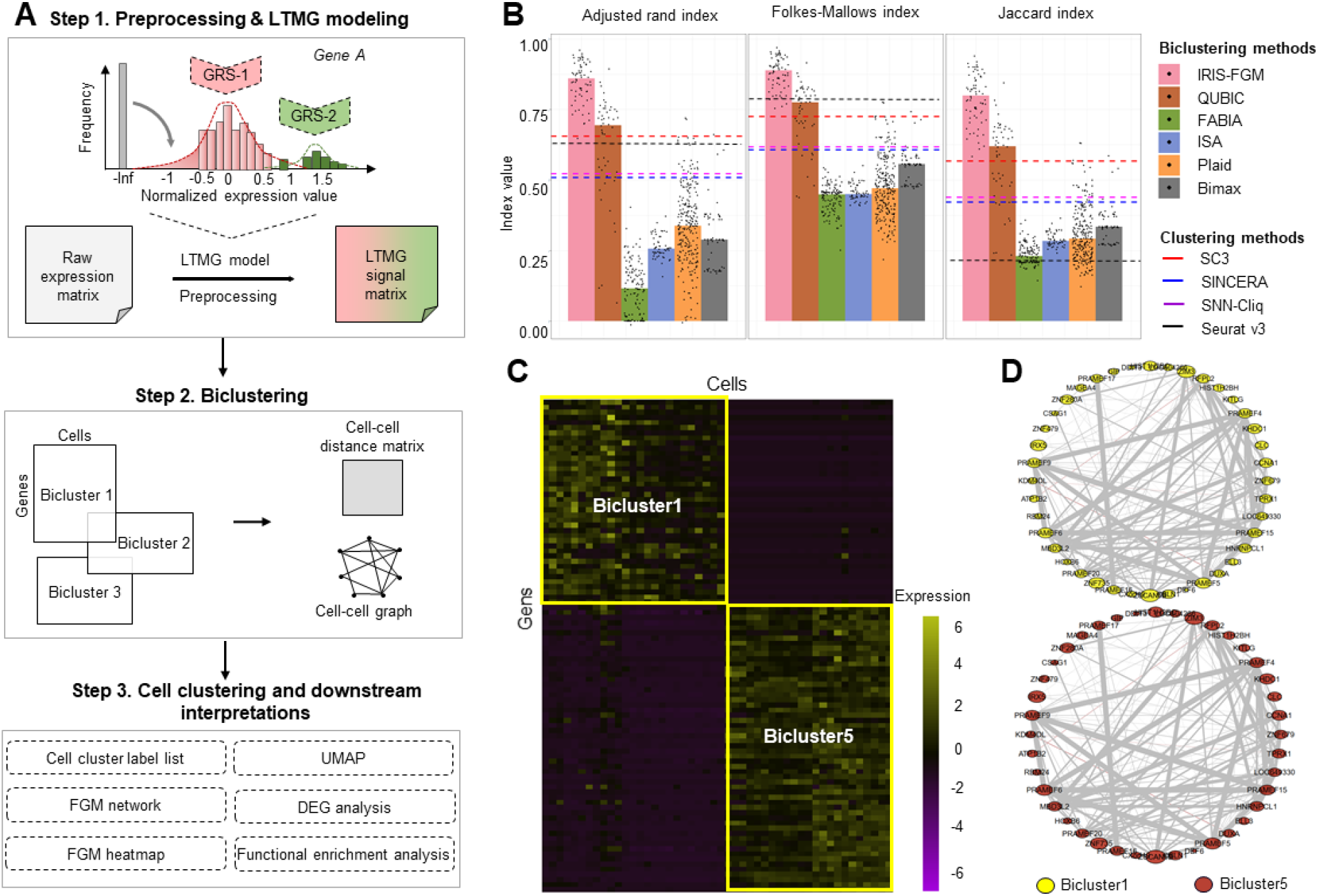
The overview of IRIS-FGM workflow and data interpretations. (A) The IRIS-FGM workflow includes three main steps: preprocessing and LTMG modeling, biclustering, and cell clustering and downstream interpretations. (B) Cell clustering evaluation, in terms of the Adjusted Rand index, Folkes-Mallows index, and Jaccard index, of IRIS-FGM against the five popular biclustering methods (bars) and three clustering methods (dashed lines) on Yan’s data. Dots represent results of different parameters used for each biclustering methods. (C) FGM heatmap visualization of two biclusters identified from Yan’s data. (D) FGM networks of Bicluster1 (yellow) and Bicluster5 (red) corresponding to C. The size of the nodes indicates the degree of presence. The thickness of edges indicates the value of the correlation coefficient. The color of the edge shows the positive (grey) and the negative (red) relationship between the two genes.

## 2 Functions and examples

IRIS-FGM contains 20 functions (more details can be found in **Supplementary Method.S2**), and the main functions of IRIS-FGM are summarized below. To further demonstrate the prowess of IRIS-FGM, we used the 90 human embryonic cells from the previous study (Yan, et al., 2013), 2700 normal human peripheral blood mononuclear cells (PBMCs) (**Supplementary Example.S1**) from 10X official website, and 1956 human CD8^+^ T cells (**Supplementary Example.S2**) from the previous study (Guo, et al., 2018) to demonstrate analysis workflow and corresponding results.

### 2.1 LTMG modeling

LTMG modeling formulates regulatory signals in each gene based on scRNA-seq data. We implement the LTMG model by function *RunLTMG* which takes the input of IRIS-FGM object and returns a discrete regulatory signal matrix. The discrete signal matrix has also been integrated into the Seurat object, which can be called by *object@LTMG@ Tmp*.*seurat*. The Seurat object with the signal matrix can be further analyzed by following its frameworks, such as cell clustering and differentially expressed gene analysis.

### 2.2 Biclustering

IRIS-FGM provides function *RunBicluster* to generate co-expressed and co-regulated gene modules depending on discretization methods, where quantile discretization methods from QUBIC 2.0 (Xie, et al., 2019) will contribute to identifying co-expression gene modules and LTMG modeling discretization method will contribute to identifying co-regulated gene modules.

### 2.3 Cell clustering and other functions

To further annotate cell heterogeneity, IRIS-FGM provides MCL-based cell clustering algorithm via function *FindClassBasedOnMC*. High accurate differentially expressed gene identification method, DEsingle (Miao, et al., 2018; Wang, et al., 2019), and pathway analysis tool, ClusterProfiler R package (Yu, et al., 2012), are integrated into IRIS-FMG to annotate cell clusters and gene signatures further. Moreover, IRIS-FMG also provides multiple visualization functions to intuitively understand cell cluster distribution and gene regulatory network (more details can be found in **Supplementary Method.S2**).

## 3 Conclusion and discussion

We developed a robust and multifunctional R package, IRIS-FGM, for scRNA-Seq data analysis that enables the identification of FGMs, cell clusters, network visualization, functional enrichment analysis of gene signatures. Furthermore, intermediate product (Seurat object with LTMG discretized matrix) of IRIS-FGM can be used for the Seurat framework. The elucidation of co-expressed and co-regulated FGMs have far-reaching impacts on how differentially activated transcriptional regulatory signals affect cell states cell, evolutionary trajectories, among other phenotypic characteristics In the long run, the knowledge derived will shed light on the computational modeling of gene regulatory network among various cell types within a complex tissue or disease microenvironment.

