## supplementary for "IRIS-FGM: an integrative single-cell RNA-Seq interpretation system for functional gene module analysis"

Supplementary Method.S1: Benchmark performance comparison regarding cell type prediction.

Supplementary Method.S2: IRIS-FGM 20 functions.

Supplementary Example.S1: Analysis based on 2700 normal human PBMCs data.

Supplementary Example.S2: Analysis based on 1956 human CD8+ T cells data.

**Supplementary Method S1. Benchmark performance comparison regarding cell type prediction.**

The Yan's scRNA-Seq dataset was used to validate the performance of biclusters-based MCL cell prediction method, compared with five biclustering methods, namely Bimax (Prelić, et al., 2006), FABIA (Hochreiter, et al., 2010), ISA (Bergmann, et al., 2003), Plaid (Ayadi, et al., 2009) and QUBIC (Li, et al., 2009), and four clustering methods, namely SC3 (Kiselev, et al., 2017), SINCERA (Guo, et al., 2015), SNN-Cliq (Xu and Su, 2015), Seurat v3 (Butler, et al., 2018). All benchmarking tools are used as the default parameter.

**Supplementary Method.S2. IRIS-FGM 20 functions**.

IRIS-FGM contains a total of 20 functions. (1) *ReadFrom10X_fold* and (2) *ReadFrom10X_h5* provide data input methods for the commercial platform 10X GENOMICS. (3) *CreateIRISFGMObject* creates an R object within the S4 data structure. (4) *AddMeta*, (5) *PlotMeta*, and (6) *SubsetData* are for calculating, visualizing the number of the detected gene, and removing low-quality cells. (7) *ProcessData* provides two methods for normalization either library size or scran normalization. (8) *RunLTMG* utilizes LTMG model to generate a LTMG signal matrix. (9) *GetLTMGmatrix* extracts the LTMG signal matrix from the object. (10) *RunDiscretization* generates the discretized matrix. (11) *RunBicluster* performs QUBIC2 biclustering method for the discretized matrix. (12) *PlotHeatmap* (13) *PlotNetwork* and (14) *PlotPlotModuleNetwork* provides visualization functions for the co-expression gene module using the qgraph R package (Epskamp, et al., 2012). (15) *FindClassBasedOnMC* uses MCL to predict cell type from identified co-expression gene modules. IRIS-FGM also integrates Seurat dimension reduction and cell clustering method and provides quick mode analysis via (16) *RundimensionReductoin* and (17)

*RunClassification*. (18) *PlotDimension* visualizes cell clusters in UMAP space. (19) *FindMarkers* identifies DEGs depended on DEsingle method (Miao, et al., 2018; Wang, et al., 2019) and (20) *RunPathway* analyzes functional pathways by implementing ClusterProfiler R package (Yu, et al., 2012).

(1) *ReadFrom10X_folder* and (2) *ReadFrom10X_h5* are used to import scRNA-Seq data from commercial platform 10X. (3) *CreateIRISFGMObject* is used to create an R object. For example:

```
> original_expression <- ReadFrom10X_h5("file.h5")
> original_expression <- ReadFrom10X_folder(“path to folder”)
> object <- CreateIRISFGMObject(original_expression)
```

(4) *AddMeta*, (5) *PlotMeta,* and (6) *SubsetData* are used to remove low quality cell. *AddMeta* can also accept customized cell label, which requires dataframe format. For example:

```
> object <- AddMeta(object, meta.info= celltype)
> PlotMeta(object)
> object <- SubsetData(object, nFeature.upper=15000,nFeature.lower=8000,
 Counts.upper=700000, Counts.lower=400000)
```

(7) *ProcessData i*s used to normalize via library size normalization or scran (optional) and impute data via DrImpute (optional). For example:

```
> object <- ProcessData(object, normalization = "LibrarySizeNormalization", IsImputation = FALSE)
```

(8) *RunLTMG* is used to generate a LTMG signal matrix, and (9) *GetLTMGmatrix* is used to obtain the LTMG signal matrix. For example:

```
> object <- RunLTMG(object, Gene_use = 500)
> LTMGmatrix <- GetLTMGmatrix(object)
```

(10) *RunDiscretization* is used to binarize the raw expression matrix. For example:

> *object <- RunDiscretization (object)*

(11) *RunBicluster* is used to perform QUBIC2 based biclustering.

> *object <- RunBicluster(object, DiscretizationModel = "Quantile",OpenDual = FALSE,*
*NumBlockOutput = 100, BlockOverlap = 0.7, BlockCellMin = 15)*

(12) *PlotHeatmap*, and (14) *PlotPlotModuleNetwork* are used to visualized FGMs. For examples:

> *PlotHeatmap(object, N.bicluster = c(1,5))*

> *PlotPlotModuleNetwork(object, N.bicluster = c(1,5))*

(13) *PlotNetwork can visualize the overlap gene or cell based on given biclusters.*

> *PlotNetwork(object, edge.by = "gene", N.bicluster = 1:20)*

(15) *FindClassBasedOnMC* predicts cell type based on MCL.

> *object <- FindClassBasedOnMC(object)*

(16) *RundimensionReduction* and (17) *RunClassification* are used for performing Seurat dimension reduction and cell clustering method. (18) *PlotDimension* can visualize Seurat result from (16) and (17). For example:

> *object <- RunDimensionReduction(object, reduction = "umap")*

> *object <- RunClassification(object, k.param = 20, resolution = 0.5, algorithm = 1)*

> *PlotDimension(object,reduction = "umap")*

(19) *FindMarkers* is used for finding DEG.

> *object <- FindMarkers(object)*

(20) *RunPathway* enables user to perform pathway enrichment analysis based on KEGG or GO for CTS marker gene.

```
> object <- RunPathway(object, selected.gene.cutoff = 0.05, species = "Human", database = "GO", genes.source = "CTS")
```

or co-expression genes in gene module.

```
> object <- RunPathway(object ,module.number = 1, selected.gene.cutoff = 0.05, species = "Human", database = "GO", genes.source = "Bicluster")
```

**Example.S1: 10X data analysis**

**S1.1 Input, preprocess, and model data**

2700 cells PMBC data set can be obtained from the 10X official website (https://support.10xgenomics.com/single-cell-gene-expression/datasets/1.1.0/pbmc3k).

(i) IRIS-FGM provides *ReadFrom10X_folder*, *ReadFrom10X_h5* to read in data from 10X platform. For example:

```
> matrix <- ReadFrom10X_folder(“./folder_10X/”)
```

(ii) *CreatIRISFGMObject* creates IRIS-FGM object. For example:

```
> object <- CreateIRISFGMObject(matrix)
Creating IRISFGM object.
The original input file contains 2700 cells and 32738 genes
Removed 16104 genes that total expression value is equal or less than 0
Removed 0 cells that number of expressed gene is equal or less than 0
```

(iii) *AddMeta* adds meta-information (cell annotation), and PlotMeta will be used for showing data quality .

```
> object <- AddMeta(object)
```

```
Do not provide meta info table for the object.
using original cell identity

> PlotMeta(object)
```

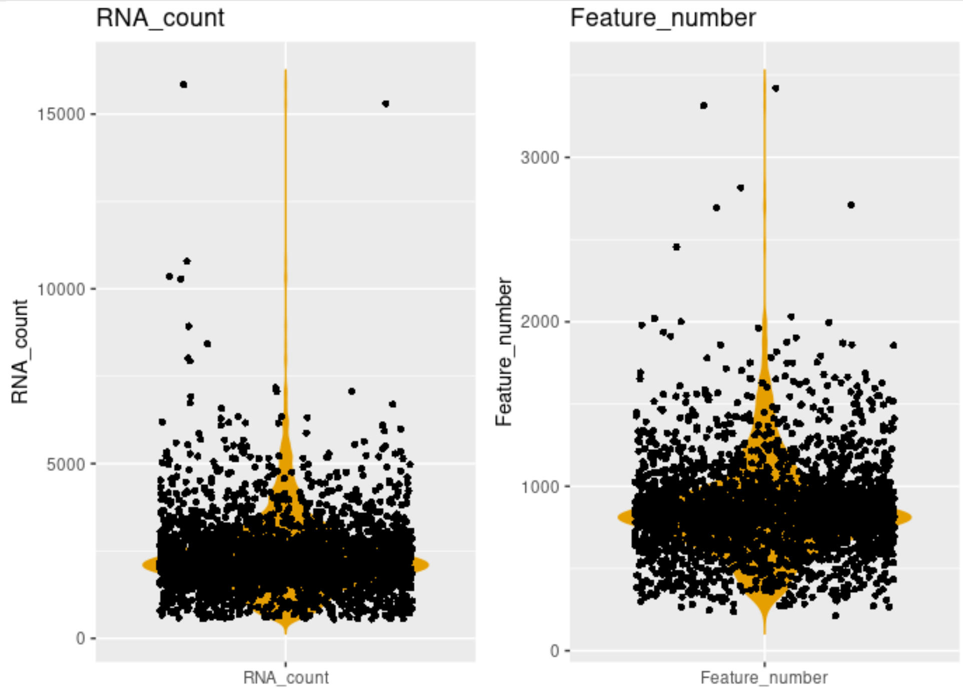


**Figure Legend:** violin plots show the distribution of the total number of RNA count (RNA_count) and the total number of genes (Feature_number).

(iv) Based on the quality figure, we can further filter out low-quality cells.

```
> object <-  SubsetData(object , nFeature.upper=3000 ,nFeature.lower=100,
                        Counts.upper= 15000, Counts.lower=1000)
```

(v) We provide two normalization methods. One is based on CPM (controlled by normalization = “LibrarSizeNormalization”). The other is based on “scan” package (controlled by normalization = “scan”).

```
> object <- ProcessData(object, normalization = "LibrarySizeNormalization")
```

### S1.2 Biclustering.

Here, we will use the biclustering algorithm to predict the co-expression gene module. In this section, the main functions are *RunDiscretization, CalBinaryMultiSignal, and RunBicluster*.

(i) Due to the large size of the data set, we will use quantile discretization which is qubic 1.0 discretization method, (Li, et al., 2009). This step can be performed by function RunDiscretization.

```
> object <- RunDiscretization (object)
writing temporary expression file ...
create temporary discretize file

QUBIC 2.1: greedy biclustering (compiled Feb 20 2020 17:11:45)

File /fs/project/PAS1475/Yuzhou_Chang/BRIC_test/10X_3K/tmp_expression.
txt contains 16634 genes by 2545 conditions
Discretization rules are written to /fs/project/PAS1475/Yuzhou_Chang/B
RIC_test/10X_3K/tmp_expression.txt.rules
Formatted data are written to /fs/project/PAS1475/Yuzhou_Chang/BRIC_te
st/10X_3K/tmp_expression.txt.chars
```

(ii) *RunBicluster* performs QUBIC 2.0 bicluster on previously created the discretized matrix. In this function, five key parameters can be controlled by the user. In this case, parameter, DiscretiationModel, can set to “quantile” for declaring we use quantile discretization; Parameter,

OpenDual, can control whether automatically extending original bicluster. If OpenDual is FALSE, the user needs to change Parameter, Extension, to decide the extension consistency level. Parameter, NumBolckOutput, controls the number of outputs biclusters. Parameter, BlockOverlap, filters out bicluster blocks, which exceeds overlap rate. For example:

```
> object <- RunBicluster(object, DiscretizationModel = "Quantile",OpenDual = FALSE,
NumBlockOutput = 1000, BlockOverlap = 0.7, BlockCellMin = 15)
QUBIC 2.1: greedy biclustering (compiled Feb 20 2020 17:11:45)

File /fs/project/PAS1475/Yuzhou_Chang/BRIC_test/10X_3K/tmp_expression.
txt.chars contains 16634 genes by 2545 conditions
Discretized data contains 3 classes with charset [ 0 1 2 ]
Formatted data are written to /fs/project/PAS1475/Yuzhou_Chang/BRIC_te
st/10X_3K/tmp_expression.txt.chars.chars
Generating seed list (minimum weight 15)
11467906 seeds generated [786.139 seconds elapsed]
Clustering started

1000 clusters are written to /fs/project/PAS1475/Yuzhou_Chang/BRIC_tes
t/10X_3K/tmp_expression.txt.chars.blocks [909.568 seconds elapsed]
```

### S1.3 FGM analysis.

In this section, IRIS-CEM provides four functions for interpreting co-expression results, including *PlotHeatmap*, *PlotNetwork, PlotModuleNetwork,* and *RunPathway*.

(i) *PlotHeatmap* can visualize two co-expression modules and show the two modules' structure; *PlotNetwork* can globally visualize relationships among identified biclusters. *PlotModuleNetwork* will generate network for showing correlation among genes within a co-expression gene module.

```
> PlotHeatmap(object,  N.bicluster =c(2,25))
```

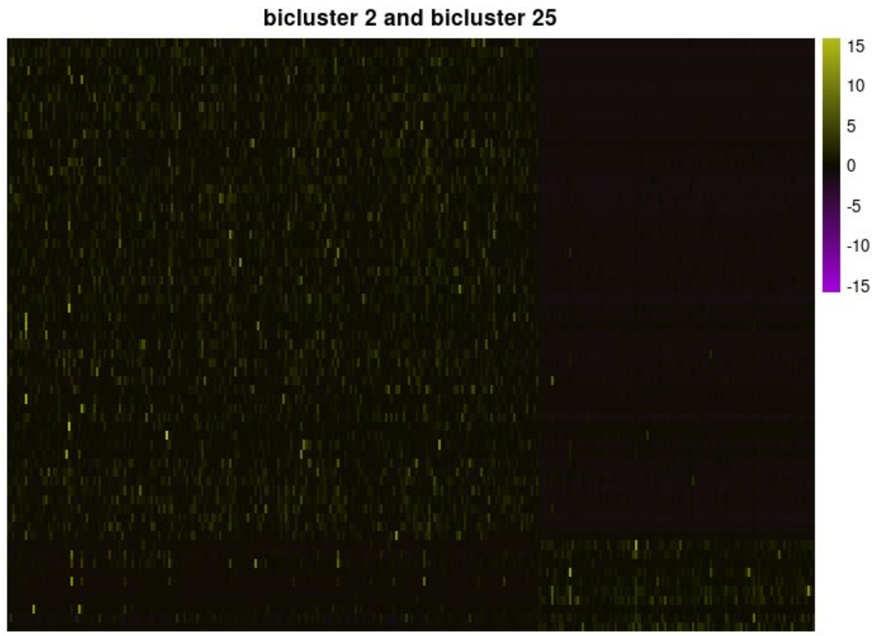


**Figure Legend:** Heatmap visualization shows the two biclusters identified in 2700 human PBMCs data, in which rows represent genes and columns represent cells. Color relates to gene express value.

(ii) PlotNetwork can show the interative of identified gene modele

```
> PlotNetwork (object, N.bicluster =c(1:5,20:26))
```

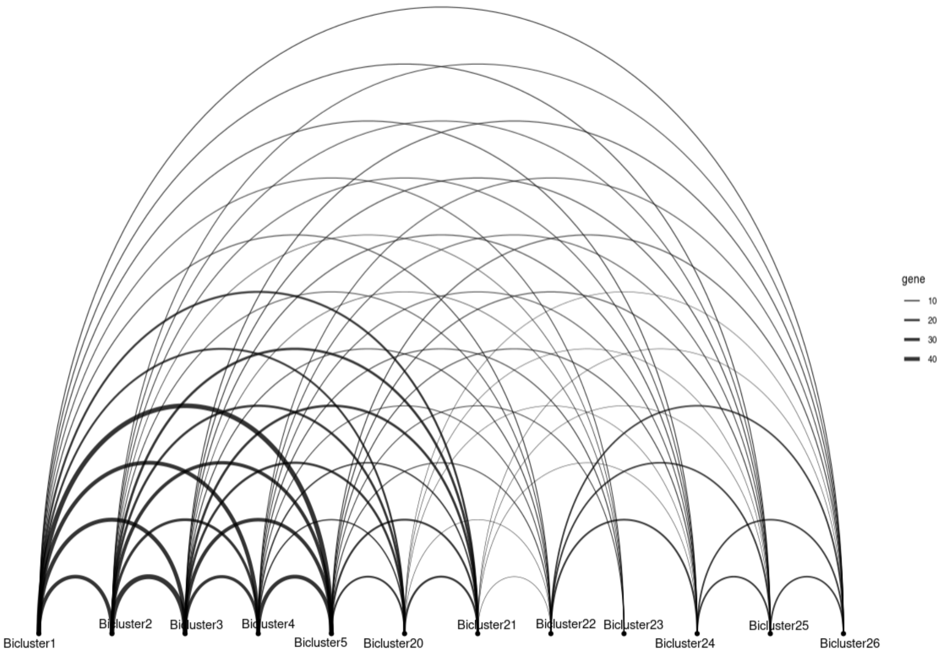


**Figure Legend:** This gene module network shows the gene overlap status among 12 gene modules. Nodes represent individual bicluster and edges represent overlapped genes. The thickness of edge means the number of overlapped gens.

(iii) Then use function *PlotModuleNetwork* shows the gene co-expression network. In this figure, gene HLA-DRB1 shows a strong positive correlation with gene HLA-DPB5 and CD74 in bicluster #1. This result means that HLA-DPB5 might potentially interact with CD74. In the Ioannis's work (Karakikes, et al., 2012), they validated HLA-DR molecules can interact with CD74.

```
> PlotModuleNetwork(object, N.bicluster = 1, Node.color = "#E8E504")
```

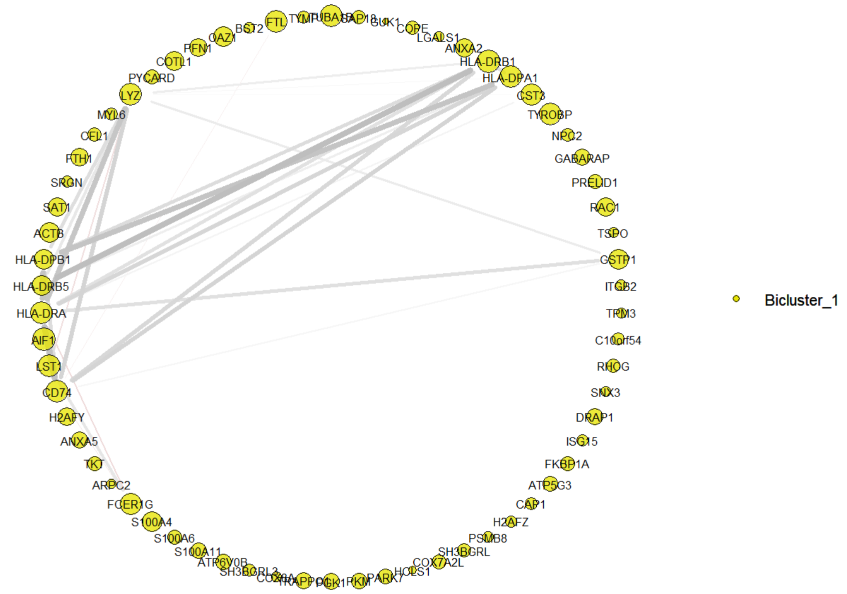


**Figure Legend**: Co-expression gene network. Yellow nodes represent the gene module network from bicluster #1. The size of the nodes indicates the degree of presence. The thickness of edges indicates the value of the correlation coefficient.

(iii) The result of functional enrichment is based on gene module (bicluster #1).

```
> object <- RunPathway(object, module.number = 1,species = "human", database = "GO", genes.source = "Bicluster", selected.gene.cutoff = 0.05)

> object@BiCluster@PathwayFromModule[1:5,]
ONTOLOGY          ID
            Description GeneRatio   BgRatio       pvalue     p.adjust
```

```
GO:0043312       BP GO:0043312
        neutrophil degranulation     19/62 485/18670 8.116792e-16 6.63
5073e-13
GO:0002283       BP GO:0002283                          neutrophil activa
tion involved in immune response     19/62 488/18670 9.084803e-16 6.63
5073e-13
GO:0042119       BP GO:0042119
           neutrophil activation     19/62 498/18670 1.315528e-15 6.63
5073e-13
GO:0002446       BP GO:0002446
    neutrophil mediated immunity     19/62 499/18670 1.364540e-15 6.63
5073e-13
GO:0002495       BP GO:0002495 antigen processing and presentation of
peptide antigen via MHC class II      8/62 101/18670 1.470752e-09 5.16
0124e-07
                 qvalue
                                                         geneID Count
GO:0043312 4.445526e-13 ANXA2/CST3/TYROBP/NPC2/RAC1/GSTP1/ITGB2/RHOG/C
AP1/PKM/TRAPPC1/S100A11/FCER1G/FTH1/LYZ/PYCARD/COTL1/BST2/FTL    19
GO:0002283 4.445526e-13 ANXA2/CST3/TYROBP/NPC2/RAC1/GSTP1/ITGB2/RHOG/C
AP1/PKM/TRAPPC1/S100A11/FCER1G/FTH1/LYZ/PYCARD/COTL1/BST2/FTL    19
GO:0042119 4.445526e-13 ANXA2/CST3/TYROBP/NPC2/RAC1/GSTP1/ITGB2/RHOG/C
AP1/PKM/TRAPPC1/S100A11/FCER1G/FTH1/LYZ/PYCARD/COTL1/BST2/FTL    19
```

```
GO:0002446 4.445526e-13 ANXA2/CST3/TYROBP/NPC2/RAC1/GSTP1/ITGB2/RHOG/C
AP1/PKM/TRAPPC1/S100A11/FCER1G/FTH1/LYZ/PYCARD/COTL1/BST2/FTL    19
GO:0002495 3.457304e-07
```

**S1.4 Seurat clustering, and visualization.**

Here we use LTMG signaling matrix as input to cluster cell and perform DEG analysis. *RunDimensionReduction* and *RunClassification* functions are for dimension reduction and cell clustering from Seurat.

(i) We need to run RunLTMG first to generate a model expression matrix. In this process, the program will automatically filter out the non-expression gene. Parameter, Gene_use, means using selected numbers of the top variant genes. For example:

```
> object <- RunLTMG (object, Gene_use = “2000”, seed =123)

number of iterations= 265
Progress:0%
number of iterations= 325
WARNING! NOT CONVERGENT!
number of iterations= 1000
WARNING! NOT CONVERGENT!
number of iterations= 1000
number of iterations= 784
number of iterations= 726
number of iterations= 910
number of iterations= 982
number of iterations= 807
```

```
Progress:10%
Progress:20%
Progress:30%
Progress:40%
Progress:50%
Progress:60%
Progress:70%
Progress:80%
Progress:90%
Progress:100%

```

(ii) Then, we use function *RunDimensionReduction* to perform dimension reduction, including PCA, t-SNE, UMAP. For example:

```
> object <- RunDimensionReduction (object, reduction= “umap”, dim = 1: 15)
Centering and scaling data matrix
  |===================================================================
======================================================================
===| 100%
Warning: The default method for RunUMAP has changed from calling Pytho
n UMAP via reticulate to the R-native UWOT using the cosine metric
To use Python UMAP via reticulate, set umap.method to 'umap-learn' and
 metric to 'correlation'
This message will be shown once per session
0%   10   20   30   40   50   60   70   80   90   100%
```

```
[----|----|----|----|----|----|----|----|----|----|
**************************************************|
```

(iii) After we get the dimension reduction result, we can use function *RunClassification* to cluster cells. The parameter, algorithm, can be selected as 1 (original Louvain algorithm), 2 (Louvain algorithm with multilevel refinement), 3 (SLM algorithm), 4 (Leiden algorithm). For example:

```
> object <- RunClassification(object, dims = 1:15, k.param = 20, resolution = 0.6,
algorithm = 1)
Computing nearest neighbor graph
Computing SNN
Modularity Optimizer version 1.3.0 by Ludo Waltman and Nees Jan van Ec
k

Number of nodes: 2545
Number of edges: 99620

Running Louvain algorithm...
0%   10   20   30   40   50   60   70   80   90   100%
[----|----|----|----|----|----|----|----|----|----|
**************************************************|
Maximum modularity in 10 random starts: 0.8575
```

```
Number of communities: 9
Elapsed time: 0 seconds
```

(iv) For showing the classification result (9 clusters), we can use function *PlotDimension* to project cell clusters on UMAP space. For example:

```
> PlotDimension(object, reduction = “umap”)

select condition to present
1 : Original
2 : ncount_RNA
3 : nFeature
4 : Seurat0.5
```

Then *PlotDimension* function will require the user to choose an identity for providing cell clusters’ information. Here we type in “4” because index “4” represents “Seurat0.5” cell clustering results.

```
> select index of cell condition: 4
```

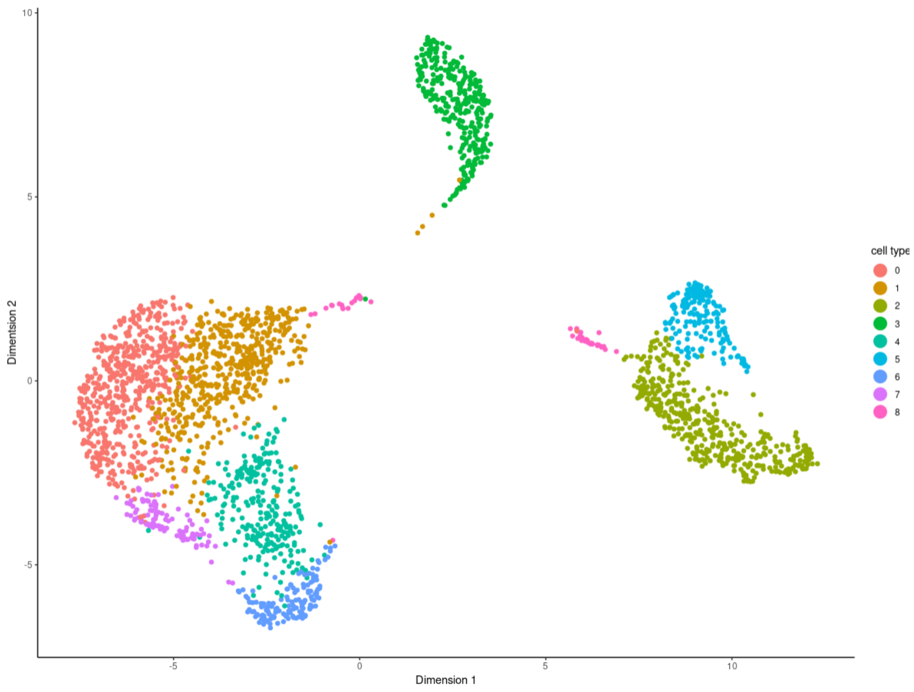


**Figure Legend:** The clustering results carried out by Seurat (9 clusters) are projected on the UMAP dimension reduction space.

**S1.5 DEG analysis.**

Here, we will use *Findmarkers* to analyze DEG based on the previous identified cell clusters. Then we utilize *RunPathway* to perform functional enrichment analysis based on identified DEG.

(i) *FindMarkers* is an interactive function and allow user to choose two groups or one group compared to all remaining unselected group. User can obtain result by using object@LTMG@MarkerGene

```
> object <- Findmarker(object)
```

Here, we have 9 cluster in total. User should first choose group with cell labe. Then user should choose an initial group and a second group (can be rest of all) to get cell-type-specific marker.

```
select condition to compare
1 : Original
2 : ncount_RNA
3 : nFeature
4 : Seurat0.5

```

We need to choose one ident as clustering label from meta information, here we use Seurat0.5 as input cell clusters.

```
select index of cell condition: 4
select index (left) of first group to compare :
1 : 0
2 : 1
3 : 2
4 : 3
5 : 4
6 : 5
7 : 6
8 : 7
9 : 8

input first group index : 1
select index (left) of second group to compare :
2 : 1
```

```
3 : 2
4 : 3
5 : 4
6 : 5
7 : 6
8 : 7
9 : 8
10 : rest of all

select index of group 2: 10
Normalizing the data
 DEsingle is analyzing 2000 of 2000 expressed genes.
```

Show the first top five results of DEGs analysis.

```
> object@LTMG@MarkerGene[1:5,]
              LFC         pval pvalue.adj.FDR
CD3D    0.2586481 1.366935e-10   2.733871e-07
RPS27   0.1457889 1.785285e-07   1.785285e-04
S100A4 -0.3020036 6.664771e-07   3.511966e-04
CCR7    0.2913331 7.023933e-07   3.511966e-04
CLIC1  -0.2999718 8.866456e-07   3.546583e-04
```

(ii) The result of pathway enrichment is based on DEGs.

```
> object <- RunPathway(object, species = “human”, database = “GO”, genes.source =
“CTS”, selected.gene.cutoff = 0.05)

> object@LTMG@Pathway[1:5,]

           ONTOLOGY         ID                      Description GeneRa
tio   BgRatio       pvalue     p.adjust       qvalue
GO:0045061       BP GO:0045061          thymic T cell selection      4
/32  21/18670 4.166703e-08 3.345862e-05 2.478092e-05
GO:0030098       BP GO:0030098       lymphocyte differentiation      8
/32 353/18670 1.068007e-07 4.288049e-05 3.175917e-05
GO:0042110       BP GO:0042110                T cell activation      8
/32 464/18670 8.530748e-07 2.076438e-04 1.537901e-04
GO:0045058       BP GO:0045058                 T cell selection      4
/32  47/18670 1.203599e-06 2.076438e-04 1.537901e-04
GO:0045059       BP GO:0045059 positive thymic T cell selection      3
/32  13/18670 1.292925e-06 2.076438e-04 1.537901e-04
                                              geneID Count
GO:0045061                       CD3D/CCR7/CD3E/CD3G     4
GO:0030098 CD3D/CCR7/CD3E/CD27/CD3G/KLF6/RHOH/SPI1     8
GO:0042110  CD3D/CCR7/CD3E/CD27/CD3G/AIF1/CD7/RHOH     8
GO:0045058                       CD3D/CCR7/CD3E/CD3G     4
```

```
GO:0045059                                   CD3D/CD3E/CD3G     3
```

**Example.S2: Immuno-oncology data**

**S2.1 download data and preprocessing**

We downloaded data (GSE99254_NSCLC.TCell.S12346.count.txt, containing 12348 cells) and from GSE99254 (Guo, et al., 2018). To better select the target cell population (CD8+ T cell in the tumor microenvironment), we also downloaded meta information from GSE99254 and save to txt file named “meta.txt.”

(i) *read.table* function for reading in the gene expression matrix.

```
> matrix <- read.table("GSE99254_NSCLC.TCell.S12346.count.txt",header = T)
```

(*ii*) Extract CD8+ T cell from matrix.

```
> my.meta <- read.delim("meta.txt",header = T)
> my.raw.rm.na <- na.omit(my.raw)
> my.raw.new <- my.raw.rm.na[,c(-1,-2)]
> rownames(my.raw.new) <- my.raw.rm.na$symbol
> # mark sure cell IDs in meta and matrix are same
> cell.id.meta <- gsub("-",".",my.meta$UniqueCell_ID)
> rownames(my.meta) <- cell.id.meta
> my.intersect.cell <- intersect(cell.id.meta,colnames(my.raw.new))
> raw.data <- my.raw.new[,my.intersect.cell]
> meta.data <- my.meta[my.intersect.cell,]
> # add sex information
```

```
> meta.data$sex <- as.character(meta.data$Patient)
> my.sex.info <- read.table("sex_info.txt",header = T,stringsAsFactors = F)
> for (i in 1:nrow(my.sex.info)){
>  tmp.patient <- my.sex.info$Patient[i]
>  tmp.sex <- my.sex.info$sex[i]
>  meta.data$sex[meta.data$sex == tmp.patient] <- tmp.sex
> }
> CD8.index <- grep("CD8", meta.data$majorCluster)
> TTC.index <- grep("TTC", meta.data$sampleType)
> CD8.TTC.index <- intersect(CD8.index,TTC.index)
> meta.data <-meta.data[CD8.TTC.index,]
> raw.data <- raw.data[,CD8.TTC.index]
```

(iii) Create IRIS-FGM object. The matrix contains 1956 CD8+ T cell.

```
> object <- CreateIRISFGMObject(matrix)
Creating IRISFGM object.
The original input file contains 1956 cells and 23370 genes
Removed 1905 genes that total expression value is equal or less than 0
Removed 0 cells that number of expressed gene is equal or less than 0
```

(iv) AddMeta adds cell annotations.

```
> object <- AddMeta(object, meta.info = meta.data)
> PlotMeta(object)
```

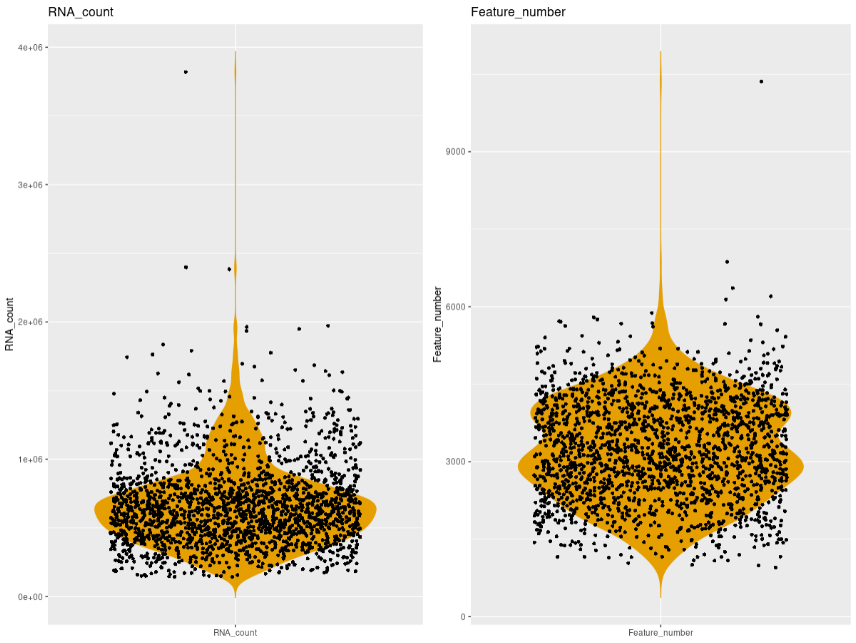


**Figure Legend:** violin plots show the distribution of the total number of RNA count (RNA_count) and the total number of genes (Feature_number).

(v) We will keep all cells in and do the normalization step.

```
> object <- ProcessData(object, normalization = "LibrarySizeNormalization",
IsImputation = FALSE, seed = 123)
```

**S2.2 Seurat clustering, and visualization**

(i) Run LTMG model on top 2000 cells:

```
> object <- RunLTMG (object, Gene_use = “2000”, seed =123)
Progress:0%
Progress:10%
Progress:20%
Progress:30%
Progress:40%
```

```
Progress:50%
Progress:60%
Progress:70%
Progress:80%
Progress:90%
Progress:100%
```

(ii) Perform dimension reduction based on Seurat:

```
> object <- RunDimensionReduction (object, reduction= “umap”, dim = 1: 15)
Centering and scaling data matrix
  |===================================================================
======================================================================
===| 100%
Warning: The default method for RunUMAP has changed from calling Pytho
n UMAP via reticulate to the R-native UWOT using the cosine metric
To use Python UMAP via reticulate, set umap.method to 'umap-learn' and
 metric to 'correlation'
This message will be shown once per session
0%   10   20   30   40   50   60   70   80   90   100%
[----|----|----|----|----|----|----|----|----|----|
**************************************************|
```

(iii) Perform cell clustering based on Seurat:

```
> object <- RunClassification(object, dims = 1:15, k.param = 20, resolution = 0.5,
algorithm = 1)
```

```
Computing nearest neighbor graph
Computing SNN
Modularity Optimizer version 1.3.0 by Ludo Waltman and Nees Jan van Ec
k

Number of nodes: 1956
Number of edges: 63146

Running Louvain algorithm...
0%   10   20   30   40   50   60   70   80   90   100%
[----|----|----|----|----|----|----|----|----|----|
**************************************************|
Maximum modularity in 10 random starts: 0.8342
Number of communities: 10
Elapsed time: 0 seconds
```

(iv) Show the clustering result on UMAP space:

```
> PlotDimension(object, reduction = “umap”)
select condition to present
1 : ncount_RNA
2 : nFeature
3 : UniqueCell_ID
4 : Patient
5 : majorCluster
```

```
6 : sampleType

7 : sex

8 : Seurat0.5
```

When R asks the user to choose an index to plot, type in the index. Here we type in “8.”

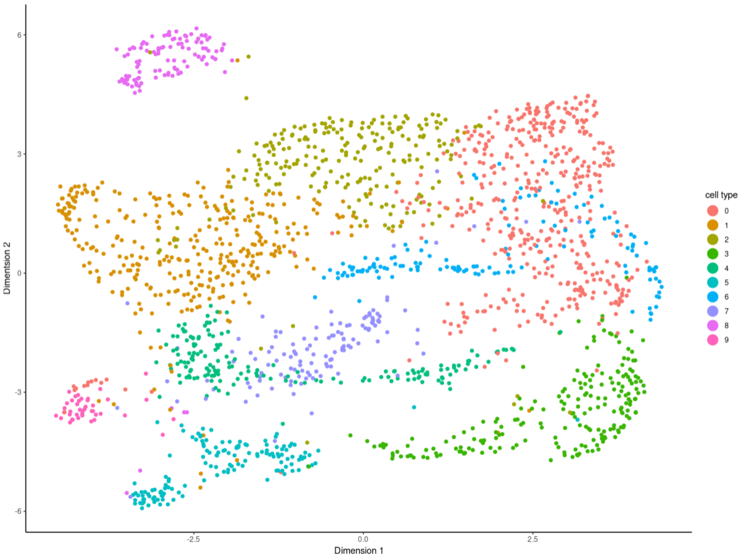


**Figure Legend:** The clustering results carried out by Seurat (9 clusters) are projected on UMAP dimension reduction space.

**S2.3 Co-expression analysis**

(i) Perform quantile discretization and biclustering.

```
> object <- RunBicluster(object = object, DiscretizationModel = “LTMG”, OpenDual =
FALSE, Extension = 1, NumBlockOutput = 100, BlockOverlap = 0.7, BlockCellMin = 15)
```

(ii) visualize bicluster information via three functions.

```
> PlotHeatmap(object,  N.block =c(1,14))
```

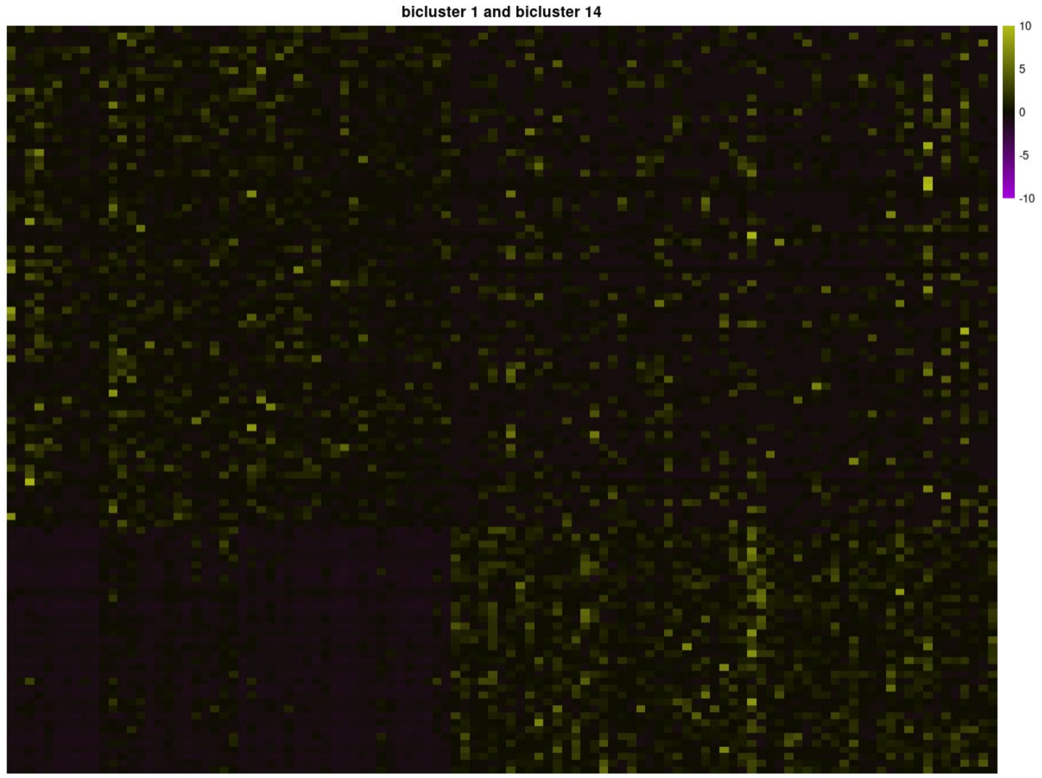


**Figure Legend:** Heatmap visualization of two biclusters identified in 1956 human CD8+ T cell in the tumor microenvironment. Rows represent genes and columns represent cells. Color relates to gene express value.

```
> PlotNetwork (object )
```

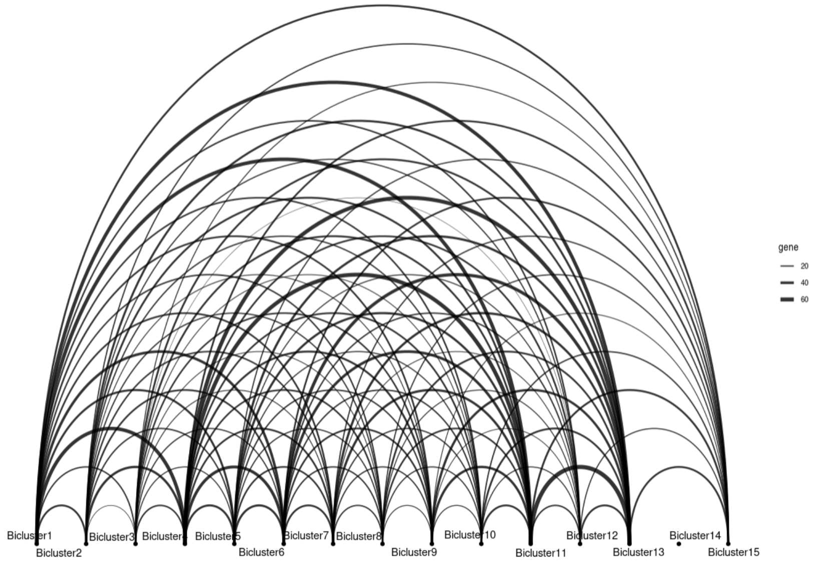


**Figure Legend:** This gene module network shows the gene overlap status among 12 gene modules. Nodes represent individual bicluster and edges represent overlapped genes. The thickness of edges means the number of overlapped genes.

```
> PlotModuleNetwork(object, N.bicluster = 1, Node.color = "#E8E504")
```

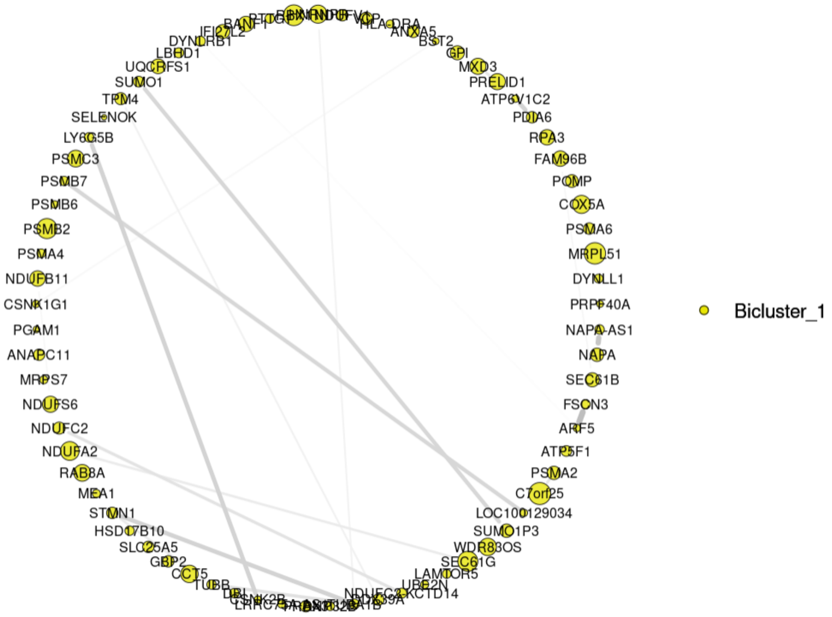

**Figure Legend**: Co-expression gene network. Yellow nodes represent the gene module network from bicluster #1. The size of the nodes indicates the degree of presence. The thickness of edges indicates the value of the correlation coefficient.

**S2.4 FGM and DEG analysis**

(i) Perform DEG analysis based on Seurat predicted cell cluster.

```
> object <- Findmarker(object)
```

Here, we will use Seurat 0.5 (index is 8) as a cell label to find DEGs regarding cluster 8.

```
select condition to compare
1 : ncount_RNA
2 : nFeature
3 : UniqueCell_ID
4 : Patient
5 : majorCluster
6 : sampleType
7 : sex
8 : Seurat0.5

select index of cell condition: 8
select index (left) of first group to compare :
1 : 0
2 : 1
3 : 2
```

```
4 : 3
5 : 4
6 : 5
7 : 6
8 : 7
9 : 8
10 : 9

input first group index : 9
select index (left) of second group to compare :
2 : 1
3 : 2
4 : 3
5 : 4
6 : 5
7 : 6
8 : 7
9 : 8
10 : 9
11 : rest of all

select index of group 2: 11
Normalizing the data
```

```
DEsingle is analyzing 2000 of 2000 expressed genes

> object@LTMG@MarkerGene[1:5,]
          LFC         pval pvalue.adj.FDR
             LFC         pval pvalue.adj.FDR
CX3CR1 0.5272759 3.319295e-07   0.0005400643
FGFBP2 0.5725258 5.400643e-07   0.0005400643
ADGRG1 0.4662232 2.421183e-06   0.0016141217
PLEK   0.4846990 1.173135e-05   0.0058656748
FCGR3A 0.5307075 8.877033e-05   0.0355081316
```

(ii) Use DEGs of cluster 1 to perform functional enrichment analysis.

```
> object <- RunPathway(object, species = “human”, database = “GO”, genes.source =
“CTS”, selected.gene.cutoff = 0.05)
> object@LTMG@Pathway[1:5,]
           ONTOLOGY         ID
        Description GeneRatio   BgRatio       pvalue    p.adjust
qvalue
GO:0070528       BP GO:0070528                                 protein
 kinase C signaling       2/5  29/18670 2.322918e-05 0.006132504 0.002
347370
GO:0008347       BP GO:0008347                                       g
lial cell migration       2/5  49/18670 6.714036e-05 0.006659824 0.002
549215
```

```
GO:2000179       BP GO:2000179 positive regulation of neural precursor
 cell proliferation       2/5  52/18670 7.567982e-05 0.006659824 0.002
549215
GO:0019932       BP GO:0019932                        second-messenger
-mediated signaling       3/5 439/18670 1.246566e-04 0.008227338 0.003
149220
GO:2000177       BP GO:2000177         regulation of neural precursor
 cell proliferation       2/5  88/18670 2.176358e-04 0.011491171 0.004
398534
                       geneID Count
GO:0070528        ADGRG1/PLEK     2
GO:0008347      CX3CR1/ADGRG1     2
GO:2000179      CX3CR1/ADGRG1     2
GO:0019932 CX3CR1/ADGRG1/PLEK     3
GO:2000177      CX3CR1/ADGRG1     2
```
